# Investigating the Role of Different Co-receptors on T-cell Activation Through IFN-γ Secretion Using Spatially Controlled Cell Monitoring Platform

**DOI:** 10.64898/2026.09.07.749865

**Authors:** Naiara Lartitegui-Meneses, Enrique Azuaje-Hualde, Andrea Merino, Sara Lopez-de-Lacalle, Fernando Benito-Lopez, Adai Colom, Lourdes Basabe-Desmonts

## Abstract

Conventional methods for studying T-cell activation typically assess receptor engagement and downstream functional outputs in separate experimental formats, thereby obscuring the spatial correlation between initial receptor arrangement and localized functional outputs. This makes it difficult to examine the individual and combined effects of receptors and co-receptors engagement within the immunological synapse within the same controlled cellular microenvironment. Building upon the recently published CellStudio platform, previously validated for monitoring growth factor interactions in adherent cell models like mesenchymal stem cells and HeLa cells, we adapted this modular system to investigate non-adherent immune cells. This platform integrates Printing and Vacuum Lithography (PnVlitho) with bead-based immunoassays to generate defined activation patterns surrounded by cytokine capture antibodies, establishing a “present-and-measure” framework for localized biosensing. Jurkat T-cells were patterned on fibronectin alone or in combination with anti-CD3 and/or anti-CD4 antibodies, enabling precise engagement and activation of the co-receptors. Cell capture, basal contact morphology, and IFN-γ secretion were evaluated for each condition. While CD4 engagement alone had a minimal effect, presenting anti-CD3 and anti-CD4 together caused the cells to spread into wide, circular contact zones and trigger the strongest local IFN-γ signals. These findings show that first adaptation of CellStudio for suspension cells enables the standardized comparison of receptor-dependent differences in Jurkat-cell capture, contact morphology, and local cytokine-associated signals within a spatially defined assay, emerging as a powerful tool for understanding co-receptor cooperation in T-cell function and immunotherapy.

## Introduction

The initiation of adaptive immune responses relies on the ability of naïve T lymphocytes to recognize specific antigens with high sensitivity and selectivity. This process, also known as immunological synapse, is mediated by the T-cell receptor (TCR), which detects peptide antigens presented by major histocompatibility complex (MHC) molecules on the surface of antigen-presenting cells.^1–3^ The TCR operates as a multichain complex in which the antigen-recognition αβ or γδ chains are physically associated with the CD3 signaling module, composed of CD3γ, CD3δ, CD3ε, and ζ chains.^4–6^ Upon antigen recognition, phosphorylation of immunoreceptor tyrosine-based activation motifs (ITAMs) within the CD3 complex initiates intracellular signaling cascades that drive T-cell activation, cytoskeletal remodeling, cytokine secretion, and immunological synapse formation. TCR signaling is further modulated by accessory receptors that tune the efficiency, duration, and functional outcome of activation. CD4 and CD8 act as co-receptors by binding MHC class II or class I molecules, respectively, and by recruiting the Src-family kinase Lck to the TCR–CD3 complex, thereby enhancing early signaling events. CD28, in contrast, functions as a co-stimulatory receptor that complements TCR-derived signals and supports transcriptional activation, survival, proliferation, and cytokine production.^7–10^ Together, these receptors organize signaling, adhesion, and actin remodeling at the immunological synapse, providing a spatially coordinated platform for T-cell activation that influence cytokine secretion.^11–15^

Despite their central role, the individual contributions of CD3-associated triggering and CD4 co-receptor engagement to T-cell activation remain incompletely defined. CD3 ligation is widely used to initiate TCR signaling experimentally, yet CD3 engagement alone does not necessarily translate into full functional activation; depending on antibody format, ligand immobilization, receptor clustering, and downstream signaling context, anti-CD3 stimulation can induce productive activation, partial signaling, or even anergic responses. ^16–19^ Moreover, receptor-proximal events do not always correlate directly with later activation outputs, as TCR downregulation and functional T-cell activation can be controlled by separable signaling mechanisms. ^20,21^ CD4, classically described as a positive amplifier of TCR sensitivity, has shown a highly context-dependent contribution to immunological synapse. In particular, CD4 may enhance antigen recognition, can have limited stabilizing effects or even inhibit helper T-cell activation depending on TCR affinity, receptor format, and activation threshold.^22–25^ These apparently divergent findings highlight that the roles of CD3 and CD4 cannot be fully understood from bulk activation readouts alone, and require experimental systems capable of isolating receptor-specific effects while preserving spatial control over cell stimulation and responses.

Dissecting the distinct roles of CD3 and CD4 remains challenging because standard analytical methods separate spatial contact mechanics from functional secretory readouts. For instance, transcriptomic approaches have been instrumental in defining activation-associated gene programs and identifying molecular signatures of T-cell stimulation. However, they provide only an indirect view of function and do not necessarily capture the timing, localization, or magnitude of secreted effector proteins.^26–28^ Conversely, cytokine-detection methods, including conventional immunoassays and emerging Raman- or plasmonic-based biosensors, directly quantify functional outputs such as IFN-γ, IL-2, or TNF. Yet, these approaches are generally optimized for analyte measurement in bulk or in formats separated from the local culture environment, limiting their ability to link cytokine secretion to defined receptor-engagement conditions and to the heterogeneity of the responding cell population. ^29–33^ As a result, receptor engagement, synapse-related cell behavior, actin remodeling, and cytokine secretion are typically measured in separate assays, averaging out cellular heterogeneity and severing the direct spatial link between the actuating antibody pattern and localized cytokine release. This methodological fragmentation limits direct comparison between stimulation conditions and contributes to the persistent uncertainty surrounding how individual receptor inputs are translated into early structural responses and downstream functional activation.

CellStudio, a platform that integrates controlled cell positioning, tunable microenvironmental stimulation, and localized secretion readouts within the same experimental architecture is well suited to address this methodological gap. The platform is based on Printing and Vacuum Lithography, which enables the reproducible generation of two-dimensional protein patterns that serve as the first functional cue controlling cell adhesion and spatial organization, surrounded by functionalized microbeads that provide sensing or signaling capabilities. ^34–36^ Its modular design allows key experimental parameters, including cell type, cluster size, adhesive geometry, protein pattern composition, bead chemistry, immobilized molecular cues, and biosensing strategy, to be independently tuned, providing a controlled framework to compare how defined microenvironmental inputs shape cell behavior. Across its previous applications, CellStudio has demonstrated compatibility with different cell models, including HeLa cells and Mesenchymal Stem cells, and with distinct bead-based detection strategies, such as immunoassays and structure-switching signaling aptamers. These implementations enabled the spatially resolved detection of secreted factors including VEGF and FGF-2 from small cell clusters, while preserving information on cell position, morphology, and cluster-to-cluster heterogeneity. The platform has also been used to modulate solid-phase presentation of bioactive molecules, showing that CellStudio can function not only as a secretion-monitoring system but also as a multifunctional tool to impose controlled biochemical cues and analyze their effects on proliferation, survival, receptor expression, morphology, and secretion within the same patterned environment.

In this study, we applied CellStudio to investigate receptor-specific activation of Jurkat T cells by using the patterned protein interface as a defined actuating cue for the presentation of anti-CD3 and anti-CD4 antibodies, either individually or in combination. While CD3/CD28 co-stimulation primarily governs secondary survival and metabolic signals, CD4 co-engagement directly dictates primary TCR-CD3 spatial assembly and proximal kinase recruitment (such as Lck) during early synapse formation. How CD4 spatial co-presentation directly modulates CD3-driven structural and secretory outputs remains poorly defined, making this combination the focus of the present study. Our strategy enabled selective stimulation of CD3- and CD4-mediated inputs across arrays of hundreds of micropatterned adhesive spots, while adapting CellStudio for the first time to non-adherent T-cells. Within this controlled microenvironment, we monitored cell adhesion and morphology, early actin cytoskeletal reorganization, and IFN-γ secretion to assess how each receptor condition contributes to synapse-related cell behavior and functional activation. By linking defined receptor engagement to spatially resolved structural and secretory outputs across hundreds of individual cell clusters within a single sample, this approach provides a robust framework for direct comparison between stimulation conditions and for dissecting the relationship between receptor presentation, actin dynamics, and cytokine-mediated T-cell activation.

**Figure 1.**
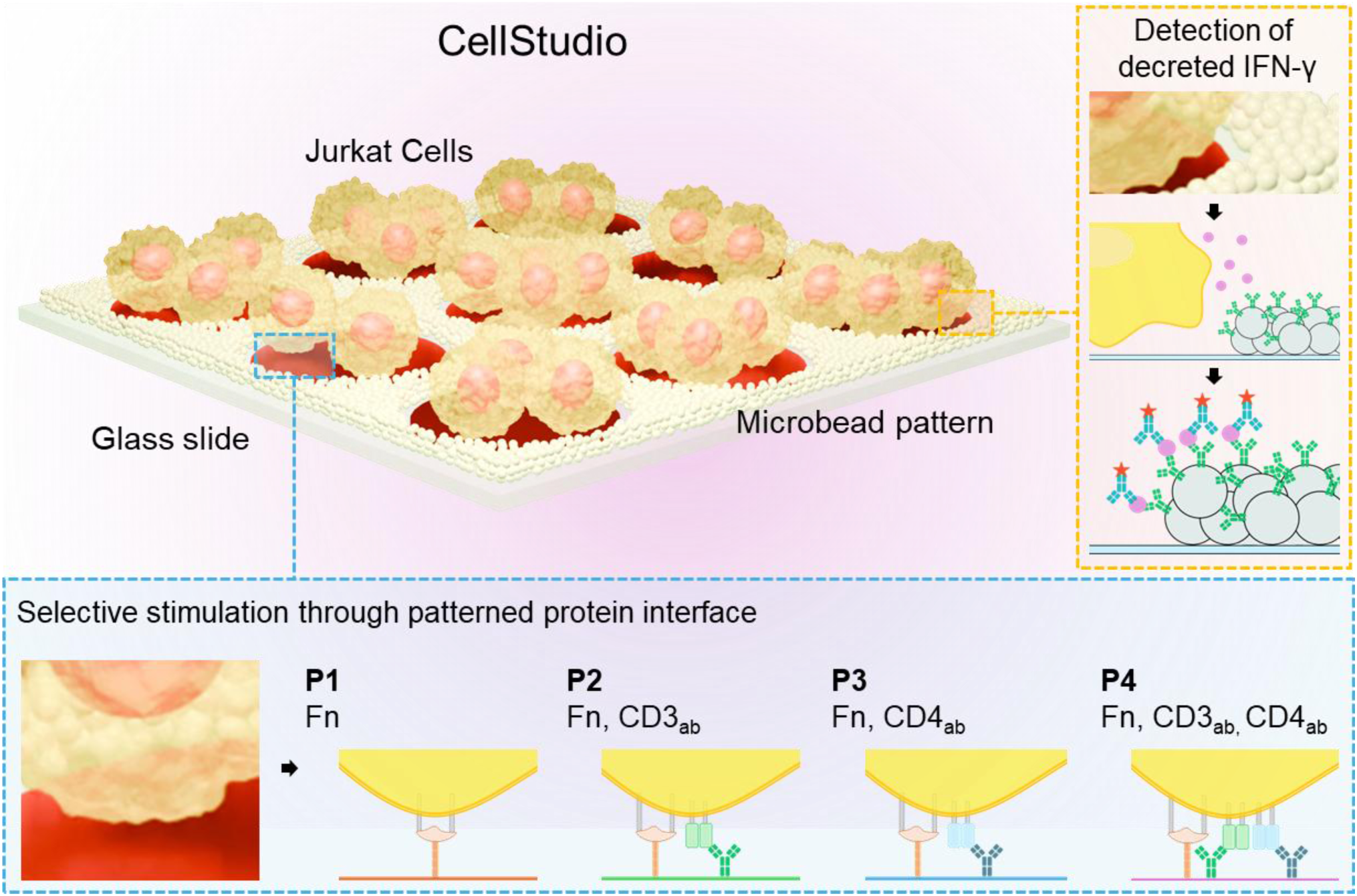
Overview of the experimental workflow. Each CellStudio substrate consists of hundreds of individual cell-attachment spots surrounded by streptavidin-coated microbeads. The composition of the spots, designed to capture and activate Jurkat cells, was varied by combining fibronectin (Fn), anti-CD3 antibodies (CD3_ab_), and anti-CD4 antibodies (CD4_ab_), enabling direct comparison of defined receptor-targeting conditions on T cell activation. In parallel with the analysis of Jurkat-cell capture and morphology, functionalization of the surrounding microbeads with anti-IFN-γ antibodies allowed localized detection, quantification, and comparison of secreted IFN-γ across the different conditions.

## 2. Experimental

### 2.1. Fabrication of CellStudio Substrates

CellStudio substrates were fabricated as previously reported, detailed in Supporting Information 1.^36^ Combined patterns of microbeads and cell-activating proteins were generated using Printing and Vacuum Lithography (PnVlitho) technique. Polydimethylsiloxane (PDMS, Ellsworth Adhesives, Spain) slabs containing channel-like structures (1000 × 5000 × 13 µm, width × length × height) with embedded micropillars (100 µm diameter, spaced 100 µm from each other) were used. Each PDMS channel was perforated to create a 2 mm inlet and a 1 mm outlet.

Four mixtures of fibronectin (Fisher Scientific, Spain) and monoclonal antibodies against CD3 (human anti-CD3 OKT3, Life Technologies, Spain) and CD4 (human anti-CD4 OKT4, Life Technologies, Spain) were prepared in 1x PBS (Sigma Aldrich, Spain), from which 4 types of micropatterned cell adhesive spots were produced, designated as Pattern 1 (P1), Pattern 2 (P2), Pattern 3 (P3), and Pattern 4 (P4). P1 consisted of fibronectin at 100 µg mL⁻¹. P2 contained fibronectin (100 µg mL⁻¹) combined with anti-CD3 OKT3 (CD3_ab_) (60 µg mL⁻¹). P3 consisted of fibronectin (100 µg mL⁻¹) mixed with anti-CD4 OTK4 (CD4_ab_) (60 µg mL⁻¹). Finally, P4 contained fibronectin (100 µg mL⁻¹) together with CD3_ab_ and CD4_ab_ (60 µg mL⁻¹ each). PDMS stamps containing micropillar arrays were incubated for 30 min with these protein solutions to allow adsorption onto the pillar surfaces. Following incubation, the slabs were rinsed with distilled water, dried under compressed air, and stamped into previously oxidized glass-bottom dishes (35 mm dish, No. 1.5 coverslip, 20 mm glass diameter, MatTek In Vitro Life Science Laboratories, Slovakia). Glass substrates were oxidized immediately prior to stamping in an air plasma chamber (29.6 W, 5 min; BlackHole, France). Transfer of the protein patterns from the PDMS to the glass substrate, allowed to generate defined activation sites for subsequent Jurkat cell adhesion and activation.

After incubation at room temperature for 15 min, the PDMS–glass assemblies were placed under vacuum (0.7 mbar, 30 min). Outlets were then sealed with tape, and 3.5 µL of streptavidin-coated polymer microbeads (CP01001, Quimigen, Spain) suspension were introduced into the inlets. Sealing prevented phase separation of the microbeads suspension during passive flow and ensured uniform distribution. Once the suspension reached the outlet, tape was removed, and assemblies were stored overnight at 4 °C to allow solvent evaporation. The following day, PDMS slabs were carefully detached from the glass, leaving the glass substrates containing microbead patterns surrounding arrays of protein activation dots.

Finally, patterned wells were blocked with 1 mL of 5% bovine serum albumin (BSA, Sigma Aldrich, Spain) in PBS 1x for 1 h at room temperature, followed by three PBS washes to remove excess blocking solution.

### 2.2. Functionalization of Microbeads

For microbeads functionalization, biotinylated IFN-γ capture antibody (BAF285, Biotechne, Spain) was diluted to 2 µg mL^-1^ in PBS and 400 µL of the suspension was added to the microbead patterns surrounding the activation sites. Substrates were incubated for 1 h at room temperature to allow biotin–streptavidin binding. The antibody solution was then removed, wells were rinsed 3 times with PBS to eliminate unbound antibody, and fresh PBS was added to maintain hydration.

### 2.3. Cell Seeding on CellStudio Substrates

Jurkat WT Clone E6-1 (ATCC) were grown in RPMI 1640 medium (Fisher Scientific, Spain) supplemented with 10% FBS (Sigma-Aldrich, Spain), 100 µg mL⁻¹ penicillin and 100 µg mL⁻¹ streptomycin at 37°C and 5% CO_2._ For patterning, cells were centrifugated and resuspend in serum-free RPMI medium (Fisher Scientific, Spain) at a concentration of 3 × 10⁵ cells mL^-1^. PBS was removed from the patterned wells, and 750 µL of the cell suspension was added in each well. Cells were incubated for 90 min at 37 °C under static conditions to promote specific adhesion to the activation dots composed of mixtures of fibronectin/CD3_ab_/CD4_ab_ (P1-P4). Cells that didn’t attach to the pattern were removed by gentle aspiration. Finally, wells were replenished with 750 µL of fresh serum free RPMI medium and incubated for 24 h to allow activation and cytokine secretion.

### 2.4. Jurkat Cell Filamentous Actin (F-actin) Staining and Basal Contact Visualization

To visualize actin and cytoskeleton re-organization upon Jurkat cells interactions with the dots, patterns (P1–P4) were generated as described above. Jurkat WT Clone E6-1 cells were centrifugated and resuspended in RPMI serum free medium for staining. 1 µM of SiR-actin (Tebubio, SC001) was added to 1mL cell suspension and incubated for 1h at 37°C and 5% CO_2_. Excess of probe was removed by centrifugation, and cells were resuspended in RPMI supplemented with 10% FBS to a final concentration of 3 × 10⁵ cells mL⁻¹. Cells were added to the CellStudio substrates and let interact with the activation dots in a fluorescence microscope at 37°C and 5% CO_2_. After 15 minutes, z-stack images of the cells were obtained. Olympus IXplore SpinSR10 microscope was used, with 60x/1.42 UPLXAPO Oil objective. Sample was excited at 640 nm, 50% power, and 500 ms of exposition time.

### 2.5. Detection of IFN-γ

IFN-γ was detected in CellStudio using a classical immunoassay, IFN-γ was captured by the biotinylated capture antibody on the beads, and then recognized by a fluorescently labeled detection antibody, forming a bead–antigen–detection antibody immunosandwich complex. To validate the immunosandwich assay for detecting cell-secreted IFN-γ, microbead patterns were prepared and functionalized as previously described, omitting the ink in the printing step. A calibration curve was generated by incubating the microbead patterns for 1 h at room temperature with recombinant human IFN-γ (PHC4031, Bio-Techne, Spain) diluted in serum-free RPMI medium at concentrations of 1, 10, and 100 ng mL⁻¹, along with a control condition without IFN-γ. After incubation, unbound protein was removed by three PBS washes. Captured IFN-γ was then detected using an Alexa Fluor 594– conjugated anti–human IFN-γ monoclonal antibody (IC2851T, Bio-Techne, Spain) incubated for 45 min at room temperature. Following three additional washes with PBS, brightfield and fluorescence images were acquired to assess signal intensity and verify sensor functionality.

For detection of IFN-γ secreted by the cells, Jurkat cells incubated in the CellStudio substrates for 24 h were fixed with 4% paraformaldehyde (Thermo Fisher, Spain) for 10 min at room temperature and washed 3 times with PBS. Secreted IFN-γ was captured by the biotinylated capture antibody immobilized on the beads and then recognized as previously explained. Samples were imaged using brightfield and fluorescence microscopy.

### 2.6. Image Acquisition and Data Analysis

Brightfield and fluorescence images were taken using an Olympus IXplore SpinSR10 spinning disk confocal super-resolution microscope equipped with a HAMAMATSU ORCA FusionBT camera, CoolLED pE-300ultra widefield light source, and filter set Ex: BP 565–585; BS: 595; Em: BP 600– 690.

Jurkat cell adhesion to the different patterns was quantified in brightfield images using ImageJ. Z-stack images of the cells were obtained using a 60x/1.42 UPLXAPO Oil objective. Basal plane of the cells was selected for segmentation using Cellpose ^37^ and manually corrected using Napari ^38^. The resulting mask was used to obtain cell shape parameters. For IFN-γ detection, fluorescence intensity was measured around each cell cluster using a ring-shaped ROI spanning the 10 µm region immediately outside the patterned dot. The measured intensity was converted into a localized IFN-γ concentration using the calibration curve. This concentration was then converted into an amount of IFN-γ using a defined analytical volume corresponding to a hemisphere with a radius of 100 µm centered on each patterned dot and was divided by to secretion time and the mean number of cells per dot for each corresponding pattern condition. Details of the process to quantify secreted IFN-γ can be found in *Supporting Information 2*.

For image visualization and preliminary analysis ImageJ was used. All statistical tests were done using GraphPad Prism 8.3

## 3. Results and Discussion

### 3.1. Experimental design

Four distinct types of CellStudio substrates designed to interact and activate different T-cell co-receptors were prepared, each containing a protein pattern in the form of dots (100 µm diameter, spaced 100 µm from each other), consisting of different mixtures of fibronectin, anti-CD3 and anti-CD4 antibodies, **Figure 2 A**. Pattern 1 (P1) consisted of fibronectin alone and served as a baseline condition, employing the limited adhesion of Jurkat cells to ECM proteins due to their low integrin activity. Pattern 2 (P2) included fibronectin mixed with anti-CD3 antibodies (CD3_ab_) to trigger T cell receptor (TCR) activation through CD3 engagement. Pattern 3 (P3) contained fibronectin with anti-CD4 antibodies (CD4_ab_) to study the isolated effects of CD4 ligation. Finally, Pattern 4 (P4) combined fibronectin, anti-CD3 (CD3_ab_), and anti-CD4 (CD4_ab_) to simultaneously stimulate both receptors and assess potential cooperative effects.

**Figure 2.**
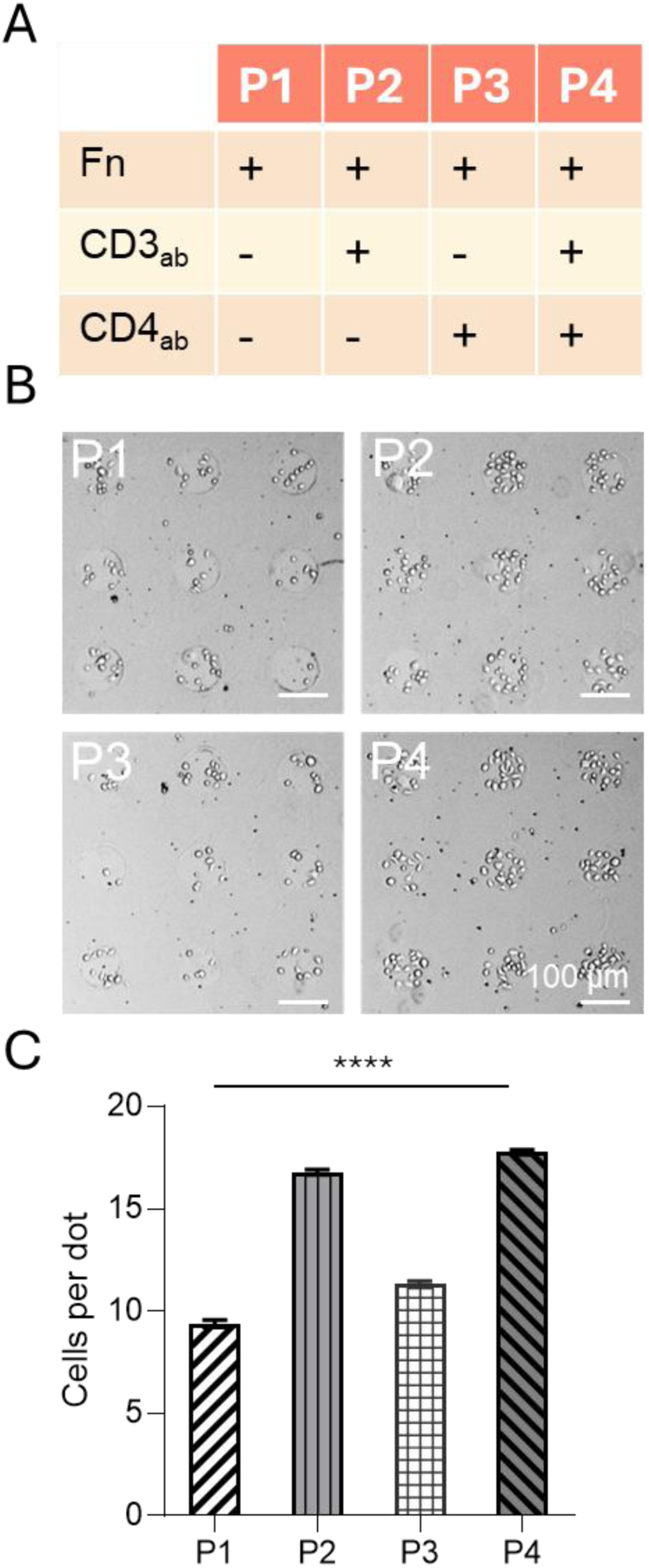
Adhesion of Jurkat T-cells to CellStudio substrates. A) Table showing the composition of the 4 types of protein patterns (P1–P4). B) Brightfield images of Jurkat T-cells attached to CellStudio substrates with the different types of protein patterns (P1–P4) after 1 h incubation. C) Average number of cells attached to each substrate (P1–P4). Individual data points represent cell counts per dot (n = 600 dots per condition across 3 independent substrates; n = 200 dots/day). Error bars represent mean ± SEM. Statistical significance was determined using the Kruskal-Wallis test (****p < 0.0001).

For all types of protein patterns, the regions surrounding each protein dot were patterned with polystyrene microbeads functionalized with anti-IFN-γ antibodies. These microbeads acted as localized biosensors, enabling fluorescence-based detection of the cytokine secretion from individual cell clusters. This integrated configuration allowed parallel assessment of Jurkat cell behavior, including adhesion, morphology, and IFN-γ secretion under precisely defined receptor stimulation conditions.

### 3.2. Analysis of CD3 and CD4 influence on adhesion of Jurkat cells on CellStudio substrates

To understand how CD3 and CD4 engagement influences early T cell recognition and physical interaction with antigen-presenting surfaces, we examined how Jurkat cells adhered to and responded morphologically to the different biochemical patterns. First, Jurkat cells were added to CellStudio substrates and incubated for 1h to allow specific cell attachment. Brightfield microscopy was used to quantify the number of Jurkat cells adhered to each 100 µm protein dot.

Adhesion levels varied depending on the biochemical composition of the protein patterns, **Figure 2 B-C**. On P1, Jurkat clusters exhibited the lowest cell counts (9 ± 4 cells per dot), which is consistent with the low affinity of T cells for fibronectin due to minimal focal adhesion machinery. Slightly increased cell capture was observed on P3 patterns containing CD4_ab_, with 11 ± 4 cells per dot, suggesting a weak interaction between T cells and the immobilized anti-CD4 antibody. In contrast, both patterns containing CD3_ab_ (P2 and P4) demonstrated the highest levels of cell attachment (16 ± 2 and 17 ± 2 cells per dot, respectively).

These results indicate that CD3 engagement enhances capture and retention of Jurkat cells with the substrate, whereas CD4 alone provides minimal additional adhesion beyond that supported by fibronectin. The similar cell count obtained from P2 and P4 suggest that CD4 engagement did not appear to further increase cell capture when CD3 is already engaged. The consistent increase in cell occupancy on CD3_ab_-containing patterns establishes CD3-targeting presentation as the principal determinant of Jurkat-cell capture in this experimental configuration. This may be a consequence of the generally higher functional expression of CD3 compared to CD4 on Jurkat cell membranes and the heterogeneity among Jurkat sublines, which can further influence surface marker abundance and adhesion behavior [29].

### 3.3. Analysis of CD3 and CD4 influence on morphology and actin dynamics of Jurkat cells on CellStudio substrates

To assess whether receptor engagement also modulated cytoskeletal dynamics and contact behavior, live-cell confocal microscopy was performed 15 minutes after cell seeding. Cells were stained with a live-cell fluorescent SiR-Actin probe and imaged at the substrate-contact plane. Morphometric parameters, including cell area and circularity, were extracted using Cellpose software ^37^ to evaluate spreading behavior and symmetry at the interface, **Figure 3**.

**Figure 3.**
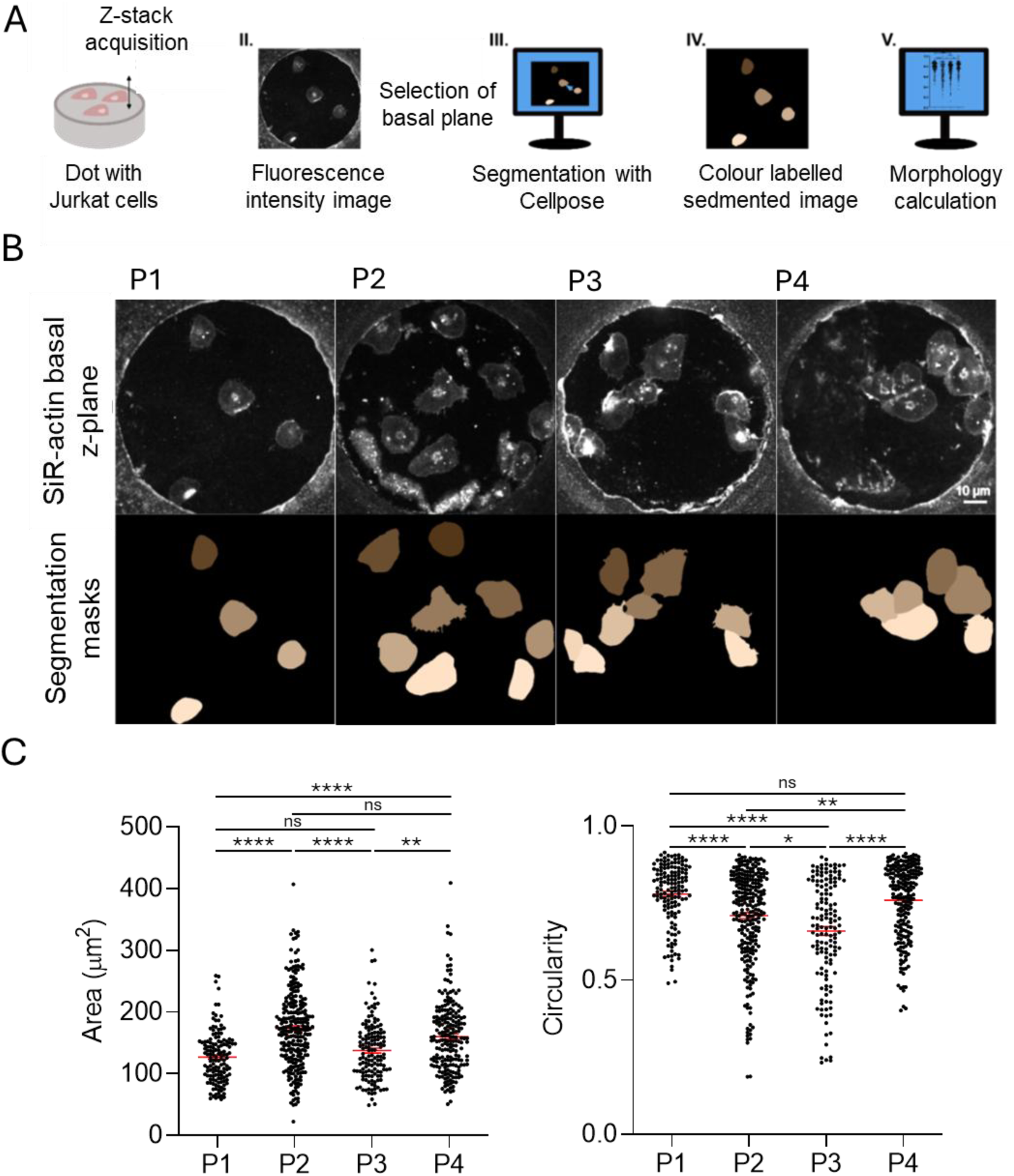
Cell morphology changes in response to different antibody-coated surfaces. A) Schematic drawing of analysis workflow. Jurkat T-cells, stained with SiR-actin, were seeded onto CellStudio substrates containing the different types of protein patterns. After 15 minutes, confocal microscopy z-stacks were acquired. The basal plane was segmented using Cellpose custom software to generate color-labeled masks for cell shape analysis. B) Top: Basal plane fluorescence images of Jurkat T-cells on proteins patterns P1-P4. Bottom: Corresponding manually curated masks. Scale bars represent 10 µm in all images C) Quantification of cell area and circularity on the 4 types of protein patterns. Each data point represents a single cell. Statistical analysis was performed using the Kruskal-Wallis test for multiple comparisons (ns. p ≥ 0.05, * p < 0.05, ** p < 0.01, **** p < 0.0001). Error bars represent mean values ± SEM.

On P1, Jurkat cells exhibited the highest circularity (0.78 ± 0.1) and the smallest contact area (127 ± 41.8 µm²), indicating limited spreading and weak fibronectin interaction. In contrast, cells on CD3_ab_-containing patterns (P2) showed pronounced morphological changes, including significant spreading and irregular contours, reflected by a reduced circularity (0.71 ± 0.15) and the largest average contact area (173.6 ± 63.5 µm²). These features are consistent with active cytoskeletal remodeling driven by T cell receptor engagement. On CD4_ab_ functionalized patterns (P3), cells demonstrated an intermediate phenotype. Circularity decreased to levels similar to P2 (0.66 ± 0.17), yet the increase in area was modest (137.5 ± 48.2 µm²), suggesting limited spreading that may reflect partial integrin modulation and weak activation associated with CD4 binding. Notably, cells on patterns containing both CD3 and CD4 (P4) displayed extensive and more symmetric spreading, with a relatively large contact area (160.4 ± 56.5 µm²) accompanied by a recovery of circular morphology (0.76 ± 0.12).

Together, these data indicate that Jurkat-cell morphology was sensitive to the receptor-targeting composition of the CellStudio patterns. CD3_ab_-containing substrates were associated with increased cell spreading. On the other hand, CD4_ab_ alone produced a more limited increase in contact area, indicating a distinct but less extensive effect of CD4 engagement in comparison to CD3. When CD3_ab_ and CD4_ab_ were presented together in P4, cells displayed a relatively large contact area together with higher circularity. The distinct response observed under combined CD3_ab_/CD4_ab_ presentation suggests that CD4 co-presentation stabilizes the interaction interface. Unlike the irregular, polarized extension seen on CD3_ab_ alone (P2), cells on combined patterns (P4) exhibit broad, isotropic spreading. This recovery in circularity (0.76 ± 0.12) alongside a large contact area 160.4 ± 56.5 µm²) reflects symmetric circumferential expansion, a structural hallmark of a stabilized, fully assembled immunological synapse rather than a lack of interaction as observed in P1.

### 3.3. Analysis of CD3 and CD4 influence on IFN-γ secretion of Jurkat cells on CellStudio substrates

While receptor engagement and morphological changes provide early indicators of T-cell activation, functional validation requires assessing cytokine secretion as a downstream output. Among the cytokines secreted by activated T lymphocytes, IFN-γ is a key effector molecule associated with immune modulation and inflammatory signaling. ^39,40^ We therefore aimed to determine how the localized activation of CD3 and CD4 influences IFN-γ secretion in Jurkat clusters patterned on CellStudio substrates.

To first validate the sensitivity of the system, we performed a calibration assay using substrates with patterned microbeads surrounding empty protein dots. These patterns were incubated with increasing concentrations of recombinant IFN-γ (0, 1, 10, and 100 ng mL⁻¹), followed by immunofluorescence labeling. Fluorescence intensity in the microbeads increased proportionally with IFN-γ concentration, indicating that the system provides good sensitivity to discern between different concentrations of the cytokine, **Figure 4**. The data were fitted to a four-parameter logistic (4PL) model. The calculated limit of detection was 0.32 ng mL⁻¹, confirming the platform’s suitability for detecting physiologically relevant cytokine levels in situ.

**Figure 4.**
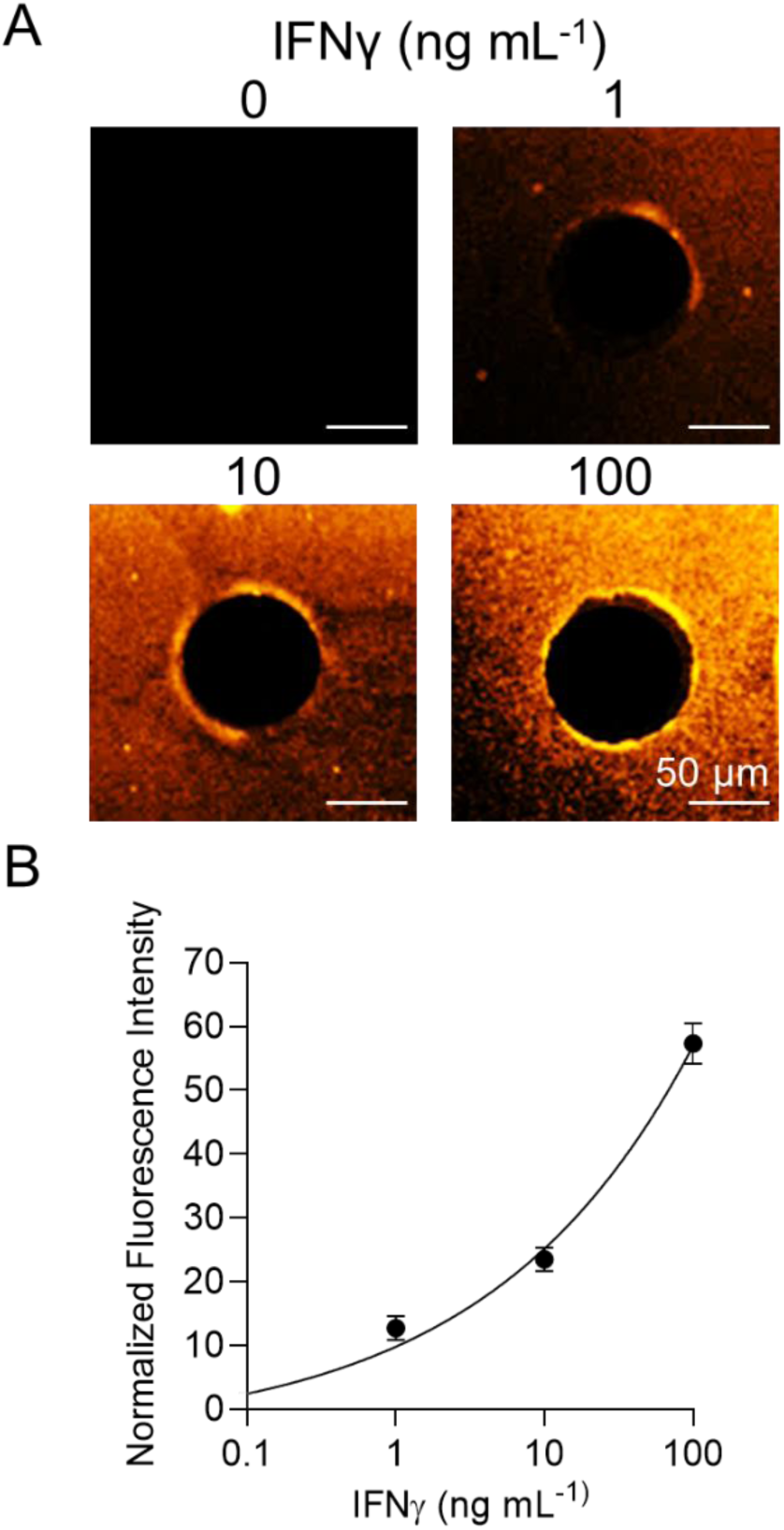
Calibration curve of IFN-γ sensor. A) Fluorescence microscopy images of CellStudio substrates with microbeads functionalized to capture IFN-γ, incubated with increasing concentrations of IFN-γ (1–100 ng mL^-1^). Fluorescence indicates binding of Alexa Fluor 596 nm anti-IFN-γ antibody. Scale bars represent 50 µm in all the images. B) Plot of the fluorescence intensity of the region of interest around each dot (10 µm annular region from the edge of the dots). Data is normalized to the mean value of the negative control (0 ng mL^-1^). Error bars represent mean values ± SEM (n = 100 dots from 3 patterns per concentration).

Following this validation, we studied IFN-γ secretion from Jurkat cells on all four types of micropatterned substrates. Jurkat cells were seeded onto the four types of biochemical patterns (P1-P4) and maintained under serum-free conditions for 24 hours to allow cytokine secretion. After incubation, IFN-γ captured by the anti-IFN-functionalized microbeads was fluorescently labeled using an immunoassay and imaged by fluorescence microscopy.

All patterns containing Jurkat clusters showed higher fluorescence intensity than the cell-free negative controls, confirming that Jurkat cells actively secreted IFN-γ under all four conditions. Minimal secretion was detected in clusters adhered to P1 (3.5 ± 0.5 ng mL^-1^ for mean localized concentration, corresponding to 0.59 ± 0.07 fg of IFN-γ per cell per day) and P3 (2.7 ± 0.3 ng mL^-1^ for mean localized concentration, corresponding to 0.6 ± 0.05 fg of IFN-γ per cell per day) patterns, suggesting that CD4 engagement alone does not robustly trigger cytokine production. In contrast, IFN-γ intensity was markedly higher in clusters on P2 (10.5 ± 1 ng mL^-1^ for mean localized concentration, corresponding to 1.5 ± 0.13 fg of IFN-γ per cell per day), indicating that CD3 activation alone is sufficient to initiate cytokine release. The highest levels of secretion were observed in the combined condition P4 (19 ± 1.5 ng mL^-1^ for mean localized concentration, corresponding to 2.71 ± 0.22 fg of IFN-γ per cell per day). This enhancement in the P4 condition suggests a synergistic effect, where CD4 co-engagement augments CD3-driven signaling pathways, resulting in more pronounced cytokine output.

**Figure 5.**
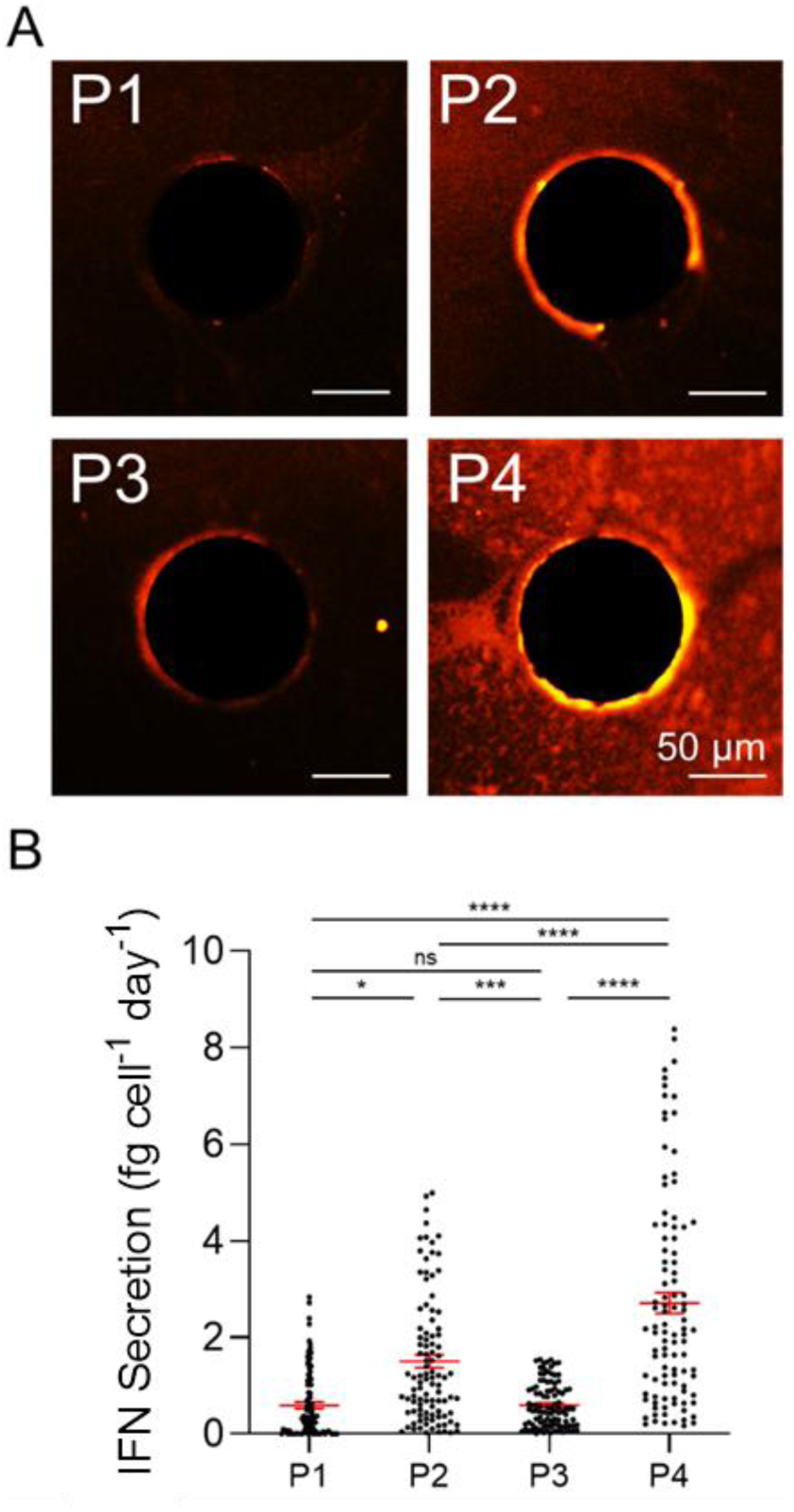
IFN-γ secretion in different co-receptor stimulation conditions. A) Fluorescence microscopy images of CellStudio substrates containing microbeads functionalized with anti-IFN-γ antibody surrounding Jurkat cell clusters in the different types of protein patterns (P1-P4) after 24 hours of incubation. Fluorescence indicates binding of Alexa Fluor 596-labelled anti-IFN-γ antibody. B) Dot-plot of the IFN-γ secretion of Jurkat cell clusters in the different types of protein patterns (P1-P4) divided by the mean number of cells per clusters in each pattern and the number of days left secreting (n = 100 cell clusters from 3 identical substrates). Each data point represents one Jurkat cell cluster. Error bars represent mean values ± SEM. Statistical significance was assessed using the non-parametric Kruskal-Wallis test (ns. p ≥ 0.05, * p < 0.05, *** p < 0.001, **** p < 0.0001).

Collectively, these results suggest that CD3 engagement is a central driver of Jurkat cell activation across multiple levels, from initial physical interaction with the substrate to the onset of functional cytokine secretion. CD3_ab_-containing patterns consistently promoted higher cell adhesion increased spreading and stronger IFN-γ output, pointing to its dominant role in initiating T cell receptor (TCR) signaling and associated cytoskeletal rearrangements. Independent CD4 engagement was associated with a more limited response profile across the morphology and secretion measurements, suggesting that it does not independently trigger robust activation under the conditions tested. However, CD4 may not act merely as a passive receptor. When presented in combination with CD3, as in the P4 condition, CD4 co-engagement appeared to enhance several aspects of Jurkat behavior, including more symmetric spreading at the contact interface and significantly elevated levels of IFN-γ secretion. The distinct response observed under combined CD3_ab_/CD4_ab_ presentation suggests that CD4 co-presentation may modulate the effects associated with CD3 engagement in this experimental configuration, potentially through mechanisms related to immunological synapse stabilization or co-receptor interactions that reinforce TCR signaling pathways.

Importantly, the integration of cell capture, contact area, circularity, and localized IFN-γ detection within a single CellStudio platform enabled these receptor-dependent effects to be compared under the same spatially controlled experimental conditions. As summarized in Table 1, each pattern generated a distinct response profile across the measured outputs, highlighting the ability of this approach to resolve condition-specific combinations of structural and functional T-cell responses.

**Table 1.** Summary response profiles for Jurkat cells on patterns P1–P4 relative to the P1 baseline. Note that the increase in circularity (+) in P4 relative to P2 reflects isotropic, symmetric synapse assembly rather than unattached cell morphology).

|  | *P1* | *P2* | *P3* | *P4* |
| --- | --- | --- | --- | --- |
| **Adhesion** | = | + | = | + |
| **Area** | = | + | = | + |
| **Circularity** | = | - | - | = |
| **IFN-γ secretion** | = | + | = | ++ |

## 4. Conclusions

In this work we investigated the specific roles of CD3 and CD4 receptors in Jurkat T cell activation. Our results show that CD3 engagement strongly drives cell adhesion and spreading, while CD4 alone induces only modest morphological changes. At the cytoskeletal level, CD3 stimulation triggers robust actin remodeling, whereas CD4 produces only partial reorganization. When both receptors are co-engaged, spreading becomes more extensive and symmetric, and actin structures are stabilized, forming a well-organized immunological synapse. Functionally, CD3 alone elicits significant IFN-γ secretion, while CD4 alone has minimal effect, indicating a low effect of CD4 on T cell activation. Combined CD3 and CD4 activation synergistically enhances cytokine release, increasing IFN-γ secretion by more than 80% compared with CD3 activation alone, demonstrating that CD4 spatial co-presentation amplifies CD3-mediated signaling by promoting symmetric, isotropic synapse contact and maximizing localized functional output. The observed differences in adhesion, morphology, and functional secretion across the four pattern types offer insight into how CD3 and CD4 may interact to shape early stages of T cell activation. Rather than acting in isolation, these receptors appear to participate in a coordinated process in which CD3 initiates activation and CD4 refines or supports the response.

Beyond these mechanistic insights, this work establishes CellStudio as a new versatile and highly accessible platform for investigating receptor-specific activation in a highly controlled spatial context. By integrating precise receptor patterning with localized cytokine sensing, the system enables a modular “present-and-measure” framework for the simultaneous assessment of adhesion, morphological dynamics, and secretory outputs, allowing detailed dissection of receptor contributions, synergistic interactions, and co-receptor modulation. Notably, this study represents the first implementation of non-adherent cells within the CellStudio platform, expanding its applicability beyond adherent models and demonstrating its versatility for suspension cell systems. Unlike more complex microsystems, CellStudiós strategic advantage lies in its simplicity; its ease of fabrication and compatibility with standard laboratory equipment significantly lower the technical barriers for high resolution cell analysis. In this sense, CellStudio provides a standardized *in vitro* framework to compare defined T cell activation scenarios under controlled spatial conditions. Future studies using primary T cells, relevant antigen-presenting systems, or antigen-specific receptor interactions could help translate this approach toward more physiologically representative models. Its adaptability and versatility can be extended to diverse biological scenarios, including alternative signaling pathways and different cell types, positioning CellStudio as a powerful bridge between receptor-level in vitro analysis and broader applications in T cell biology, immune cell engineering and immunotherapy.

## Author contributions

N. L-M., E.A.-H.: methodology, investigation, writing─review and editing, visualization. S.L-d-C., A.M.: methodology, investigation, writing─review. F.B.-L., A.C., A.A.: conceptualization, methodology, writing─review and editing, visualization, resources, supervision, project administration, funding acquisition. L.B.-D.: conceptualization, investigation, methodology, writing─review and editing, visualization, resources, supervision, project administration, funding acquisition. All authors have given approval to the final version of the manuscript.

## Data Availability

The data that support the findings of this study are available from the corresponding author, L.B-D., upon request.

## Acknowledgements

This work was funded by the Ministerio de Ciencia, Innovación y Universidades, the Agencia Estatal de Investigación (AEI, 10.13039/501100011033), and the European Regional Development Fund (FEDER, EU), under project PID2024-155781NB-I00 and from Basque Government, under Grupos Consolidados with Grant No. IT1971-26 and ELKARTEK project: K-2026/00026. EAH acknowledges funding from Basque Government Postdoctoral Program under grant POS2023_1_0014. AM acknowledges funding support from Basque Government Predoctoral Program. The authors also thank for technical support provided by “Analytical and High-Resolution Microscopy in Biomedicine” SGIker (UPV/EHU/ ERDF, EU). The authors gratefully acknowledge the Basque Resource for Advanced Light Microscopy (BRALM) located at Instituto Biofisika (CSIC, UPV/EHU) for their support and assistance in this work. We sincerely thank Prof. Alicia Alonso for her invaluable support in fostering the collaboration between our research groups and facilitating the resources that made this work possible.

## References

(1) Dustin, M. L.; Choudhuri, K. Signaling and Polarized Communication Across the T Cell Immunological Synapse. Annu. Rev. Cell Dev. Biol. 2016, 32 (1), 303–325. 10.1146/annurev-cellbio-100814-125330.

(2) Shah, K.; Al-Haidari, A.; Sun, J.; Kazi, J. U. T Cell Receptor (TCR) Signaling in Health and Disease. Signal Transduct. Target. Ther. 2021, 6 (1), 412. 10.1038/s41392-021-00823-w.

(3) Mariuzza, R. A.; Agnihotri, P.; Orban, J. The Structural Basis of T-Cell Receptor (TCR) Activation: An Enduring Enigma. Journal of Biological Chemistry 2020, 295 (4), 914–925. 10.1074/jbc.REV119.009411.

(4) Menon, A. P.; Moreno, B.; Meraviglia-Crivelli, D.; Nonatelli, F.; Villanueva, H.; Barainka, M.; Zheleva, A.; van Santen, H. M.; Pastor, F. Modulating T Cell Responses by Targeting CD3. Cancers (Basel). 2023, 15, 1189. 10.3390/cancers15041189.

(5) Hoque, M.; Grigg, J. B.; Ramlall, T.; Jones, J.; McGoldrick, L. L.; Lin, J. C.; Olson, W. C.; Smith, E.; Franklin, M. C.; Zhang, T.; Saotome, K. Structural Characterization of Two Γδ TCR/CD3 Complexes. Nat. Commun. 2025, 16 (1), 318. 10.1038/s41467-024-55467-5.

(6) Saotome, K.; Dudgeon, D.; Colotti, K.; Moore, M. J.; Jones, J.; Zhou, Y.; Rafique, A.; Yancopoulos, G. D.; Murphy, A. J.; Lin, J. C.; Olson, W. C.; Franklin, M. C. Structural Analysis of Cancer-Relevant TCR-CD3 and Peptide-MHC Complexes by CryoEM. Nat. Commun. 2023, 14 (1), 2401. 10.1038/s41467-023-37532-7.

(7) Onnis, A.; Baldari, C. T. Orchestration of Immunological Synapse Assembly by Vesicular Trafficking. Front. Cell Dev. Biol. 2019, 7. 10.3389/fcell.2019.00110.

(8) Qin, Z.; Xu, T. Deciphering the Deterministic Role of TCR Signaling in T Cell Fate Determination. Front. Immunol. 2025, 16. 10.3389/fimmu.2025.1562248.

(9) Capitani, N.; Baldari, C. T. The Immunological Synapse: An Emerging Target for Immune Evasion by Bacterial Pathogens. Front. Immunol. 2022, 13. 10.3389/fimmu.2022.943344.

(10) Thauland, T. J.; Hu, K. H.; Bruce, M. A.; Butte, M. J. Cytoskeletal Adaptivity Regulates T Cell Receptor Signaling. Sci. Signal. 2017, 10 (469). 10.1126/scisignal.aah3737.

(11) Rodrigues, L. S.; Barreto, A. S.; Bomfim, L. G. S.; Gomes, M. C.; Ferreira, N. L. C.; da Cruz, G. S.; Magalhães, L. S.; de Jesus, A. R.; Palatnik-de-Sousa, C. B.; Corrêa, C. B.; de Almeida, R. P. Multifunctional, TNF-α and IFN-γ-Secreting CD4 and CD8 T Cells and CD8High T Cells Are Associated With the Cure of Human Visceral Leishmaniasis. Front. Immunol. 2021, 12. 10.3389/fimmu.2021.773983.

(12) Chao, Z.; Mei, Q.; Yang, C.; Luo, J.; Liu, P.; Peng, H.; Guo, X.; Yin, Z.; Li, L.; Wang, Z. Immunological Synapse: Structures, Molecular Mechanisms and Therapeutic Implications in Disease. Signal Transduct. Target. Ther. 2025, 10 (1), 254. 10.1038/s41392-025-02332-6.

(13) Wipa, P.; Paensuwan, P.; Ngoenkam, J.; Woessner, N. M.; Minguet, S.; Schamel, W. W.; Pongcharoen, S. Actin Polymerization Regulates Recruitment of Nck to CD3 *ε* upon T-cell Receptor Triggering. Immunology 2020, 159 (3), 298–308. 10.1111/imm.13146.

(14) Lam, T. T.; Chong, M. M. W. Regulation of Actin Cytoskeletal Dynamics in T Cell Development and Function. Front. Immunol. 2025, 16. 10.3389/fimmu.2025.1622928.

(15) Sun, L.; Su, Y.; Jiao, A.; Wang, X.; Zhang, B. T Cells in Health and Disease. Signal Transduct. Target. Ther. 2023, 8 (1), 235. 10.1038/s41392-023-01471-y.

(16) Tsopoulidis, N.; Kaw, S.; Laketa, V.; Kutscheidt, S.; Baarlink, C.; Stolp, B.; Grosse, R.; Fackler, O. T. T Cell Receptor–Triggered Nuclear Actin Network Formation Drives CD4 ^+^ T Cell Effector Functions. Sci. Immunol. 2019, 4 (31). 10.1126/sciimmunol.aav1987.

(17) Smith, J. A.; Tso, J. Y.; Clark, M. R.; Cole, M. S.; Bluestone, J. A. Nonmitogenic Anti-CD3 Monoclonal Antibodies Deliver a Partial T Cell Receptor Signal and Induce Clonal Anergy. J. Exp. Med. 1997, 185 (8), 1413–1422. 10.1084/jem.185.8.1413.

(18) Wolf, H.; Müller, Y.; Salmen, S.; Wilmanns, W.; Jung, G. Induction of Anergy in Resting Human T Lymphocytes by Immobilized Anti-CD3 Antibodies. Eur. J. Immunol. 1994, 24 (6), 1410–1417. 10.1002/eji.1830240626.

(19) Jassin, M.; E Silva, B.; Ormenese, S.; Baron, F.; Ehx, G.; Caers, J. Comparable Restimulation of Human T Cells Activated with CD3/CD28 Beads versus Soluble Antibody Complexes. Sci. Rep. 2026, 16 (1), 9739. 10.1038/s41598-026-43542-4.

(20) van der Donk, L. E. H.; Ates, L. S.; van der Spek, J.; Tukker, L. M.; Geijtenbeek, T. B. H.; van Heijst, J. W. J. Separate Signaling Events Control TCR Downregulation and T Cell Activation in Primary Human T Cells. Immun. Inflamm. Dis. 2021, 9 (1), 223–238. 10.1002/iid3.383.

(21) Johnson, D. K.; Magoffin, W.; Myers, S. J.; Finnell, J. G.; Hancock, J. C.; Orton, T. S.; Persaud, S. P.; Christensen, K. A.; Weber, K. S. CD4 Inhibits Helper T Cell Activation at Lower Affinity Threshold for Full-Length T Cell Receptors Than Single Chain Signaling Constructs. Front. Immunol. 2021, 11. 10.3389/fimmu.2020.561889.

(22) Mørch, A. M.; Bálint, Š.; Santos, A. M.; Davis, S. J.; Dustin, M. L. Coreceptors and TCR Signaling – the Strong and the Weak of It. Front. Cell Dev. Biol. 2020, 8. 10.3389/fcell.2020.597627.

(23) Rushdi, M. N.; Pan, V.; Li, K.; Choi, H.-K.; Travaglino, S.; Hong, J.; Griffitts, F.; Agnihotri, P.; Mariuzza, R. A.; Ke, Y.; Zhu, C. Cooperative Binding of T Cell Receptor and CD4 to Peptide-MHC Enhances Antigen Sensitivity. Nat. Commun. 2022, 13 (1), 7055. 10.1038/s41467-022-34587-w.

(24) Horkova, V.; Drobek, A.; Paprckova, D.; Niederlova, V.; Prasai, A.; Uleri, V.; Glatzova, D.; Kraller, M.; Cesnekova, M.; Janusova, S.; Salyova, E.; Tsyklauri, O.; Kadlecek, T. A.; Krizova, K.; Platzer, R.; Schober, K.; Busch, D. H.; Weiss, A.; Huppa, J. B.; Stepanek, O. Unique Roles of Co-Receptor-Bound LCK in Helper and Cytotoxic T Cells. Nat. Immunol. 2023, 24 (1), 174–185. 10.1038/s41590-022-01366-0.

(25) Morales-Martínez, M.; Andón-García, D.; Patiño-Santiago, K. A.; Parga-Ortega, J. M.; Hernández-Hernández, A.; Aquino-Jarquin, G.; Patino-Lopez, G. Identification of Potential New T Cell Activation Molecules: A Bioinformatic Approach. Sci. Rep. 2024, 14 (1), 22219. 10.1038/s41598-024-73003-9.

(26) Rade, M.; Böhlen, S.; Neuhaus, V.; Löffler, D.; Blumert, C.; Merz, M.; Köhl, U.; Dehmel, S.; Sewald, K.; Reiche, K. A Time-Resolved Meta-Analysis of Consensus Gene Expression Profiles during Human T-Cell Activation. Genome Biol. 2023, 24 (1), 287. 10.1186/s13059-023-03120-7.

(27) Schmidt, R.; Steinhart, Z.; Layeghi, M.; Freimer, J. W.; Bueno, R.; Nguyen, V. Q.; Blaeschke, F.; Ye, C. J.; Marson, A. CRISPR Activation and Interference Screens Decode Stimulation Responses in Primary Human T Cells. Science (1979). 2022, 375 (6580). 10.1126/science.abj4008.

(28) Shimizu, J.; Sasaki, T.; Ong, G. H.; Koketsu, R.; Samune, Y.; Nakayama, E. E.; Nagamoto, T.; Yamamoto, Y.; Miyazaki, K.; Shioda, T. IFN-γ Derived from Activated Human CD4+ T Cells Inhibits the Replication of SARS-CoV-2 Depending on Cell-Type and Viral Strain. Sci. Rep. 2024, 14 (1), 26660. 10.1038/s41598-024-77969-4.

(29) Platchek, M.; Lu, Q.; Tran, H.; Xie, W. Comparative Analysis of Multiple Immunoassays for Cytokine Profiling in Drug Discovery. SLAS Discovery 2020, 25 (10), 1197–1213. 10.1177/2472555220954389.

(30) Chen, H.; Zhang, H.; Duan, L.; Sun, K.; Li, R.; Wang, Y.; Yao, J.; Yan, C.; Liu, Y.; Wu, Y.; Yang, G.; Han, C. Ultrasensitive SERS Detection of Cytokines through Specific Binding and Multiple Reporter Molecules. Anal. Chim. Acta 2025, 1369, 344355. 10.1016/j.aca.2025.344355.

(31) He, J.; Zhou, L.; Huang, G.; Shen, J.; Chen, W.; Wang, C.; Kim, A.; Zhang, Z.; Cheng, W.; Dai, S.; Ding, F.; Chen, P. Enhanced Label-Free Nanoplasmonic Cytokine Detection in SARS-CoV-2 Induced Inflammation Using Rationally Designed Peptide Aptamer. ACS Appl. Mater. Interfaces 2022, 14 (43), 48464–48475. 10.1021/acsami.2c14748.

(32) Baek, S. H.; Song, H. W.; Lee, S.; Kim, J.-E.; Kim, Y. H.; Wi, J.-S.; Ok, J. G.; Park, J. S.; Hong, S.; Kwak, M. K.; Lee, H. J.; Nam, S.-W. Gold Nanoparticle-Enhanced and Roll-to-Roll Nanoimprinted LSPR Platform for Detecting Interleukin-10. Front. Chem. 2020, 8. 10.3389/fchem.2020.00285.

(33) Azuaje-Hualde, E.; Lartitegi, N.; Benito-Lopez, F.; Basabe-Desmonts, L. Non-Invasive Detection of VEGF Secretion from Small Clusters of Mesenchymal Stem Cells Using VEGF-SSSA Integrated into the CellStudio Platform. Microchimica Acta 2025, 192 (8), 484. 10.1007/s00604-025-07319-2.

(34) Azuaje-Hualde, E.; Lartitegui-Meneses, N.; Alonso-Cabrera, J.; Alvarez-Braña, Y.; Martínez de Pancorbo, M.; Benito-Lopez, F.; Basabe-Desmonts, L. Analyzing the Relationship between Solid-Phase Molecular Presentation and Cell Proliferation, Morphology and Secretion Using CellStudio. ACS Appl. Mater. Interfaces 2026, 18 (4), 6423–6432. 10.1021/acsami.5c18270.

(35) Azuaje-Hualde, E.; Lartitegui-Meneses, N.; Alonso-Cabrera, J.; Inchaurraga-Llamas, A.; Alvarez-Braña, Y.; Martínez-dePancorbo, M.; Benito-Lopez, F.; Basabe-Desmonts, L. CellStudio: A Modular, Tunable and Accessible Platform for Analysis of Growth Factors Secretions in Cell Cultures. ACS Appl. Mater. Interfaces 2025, 17 (6), 8914–8923. 10.1021/acsami.4c17189.

(36) Stringer, C.; Wang, T.; Michaelos, M.; Pachitariu, M. Cellpose: A Generalist Algorithm for Cellular Segmentation. Nat. Methods 2021, 18 (1), 100–106. 10.1038/s41592-020-01018-x.

(37) Selzer, G. J.; Rueden, C. T.; Hiner, M. C.; Evans, E. L.; Harrington, K. I. S.; Eliceiri, K. W. Napari-Imagej: ImageJ Ecosystem Access from Napari. Nat. Methods 2023, 20 (10), 1443–1444. 10.1038/s41592-023-01990-0.

(38) Casanova, J.-L.; MacMicking, J. D.; Nathan, C. F. Interferon-**γ** and Infectious Diseases: Lessons and Prospects. Science (1979). 2024, 384 (6693). 10.1126/science.adl2016.

(39) Han, J.; Wu, M.; Liu, Z. Dysregulation in IFN-γ Signaling and Response: The Barricade to Tumor Immunotherapy. Front. Immunol. 2023, 14. 10.3389/fimmu.2023.1190333.

