## Supporting Information for "Investigating the Role of Different Co-receptors on T-cell Activation Through IFN-γ Secretion Using Spatially Controlled Cell Monitoring Platform"

### Supporting Information 1 : Fabrication of CellStudio Substrates by Printing and Vacuum Lithography

CellStudio substrates were generated using Printing and Vacuum Lithography (PnVLitho), a fabrication strategy that combines microcontact printing with vacuum-assisted deposition to produce spatially organized patterns of protein spots and microbeads on glass surfaces, **Figure SI-1**. In the first step, microstructured PDMS stamps containing arrays of cylindrical pillars were incubated with the corresponding protein mixtures and brought into contact with plasma-treated glass substrates. This enabled the dry transfer of circular protein spots with the same diameter as the PDMS pillars. In the present study, these spots were composed of fibronectin alone or fibronectin combined with anti-CD3 and/or anti-CD4 antibodies, thereby defining the regions for Jurkat-cell capture and receptor-specific stimulation. The same PDMS stamp guided the deposition of microbeads around the printed spots. This step relies on the gas permeability of PDMS. When the assembled PDMS–glass device is placed under vacuum, air is removed from the polymer. Once atmospheric pressure is restored, the PDMS generates a negative pressure that drives the flow of a microbead suspension through the spaces between the pillars. As a result, microbeads are deposited around each printed protein spot while leaving the central cell-attachment area accessible. Following solvent evaporation and removal of the PDMS stamp, the final CellStudio substrate consists of arrays of protein spots surrounded by patterned microbeads.

The substrates were then blocked with a 5% BSA solution in PBS for 1 h at room temperature to reduce nonspecific cell attachment and protein adsorption. Jurkat cells were added to the blocked substrates and maintained for 1 h at 37 °C and 5% $CO_2$ to allow cell capture on the patterned spots. After incubation, the supernatant containing unbound cells was removed, and fresh medium was added. In this work, the microbeads were functionalized with anti-IFN-γ capture antibodies, allowing localized cytokine detection around each Jurkat-cell cluster. This configuration enabled the generation of hundreds of spatially separated presenting units in which cell capture, morphology, and IFN-γ-associated signal could be analyzed under defined CD3/CD4 presentation conditions.

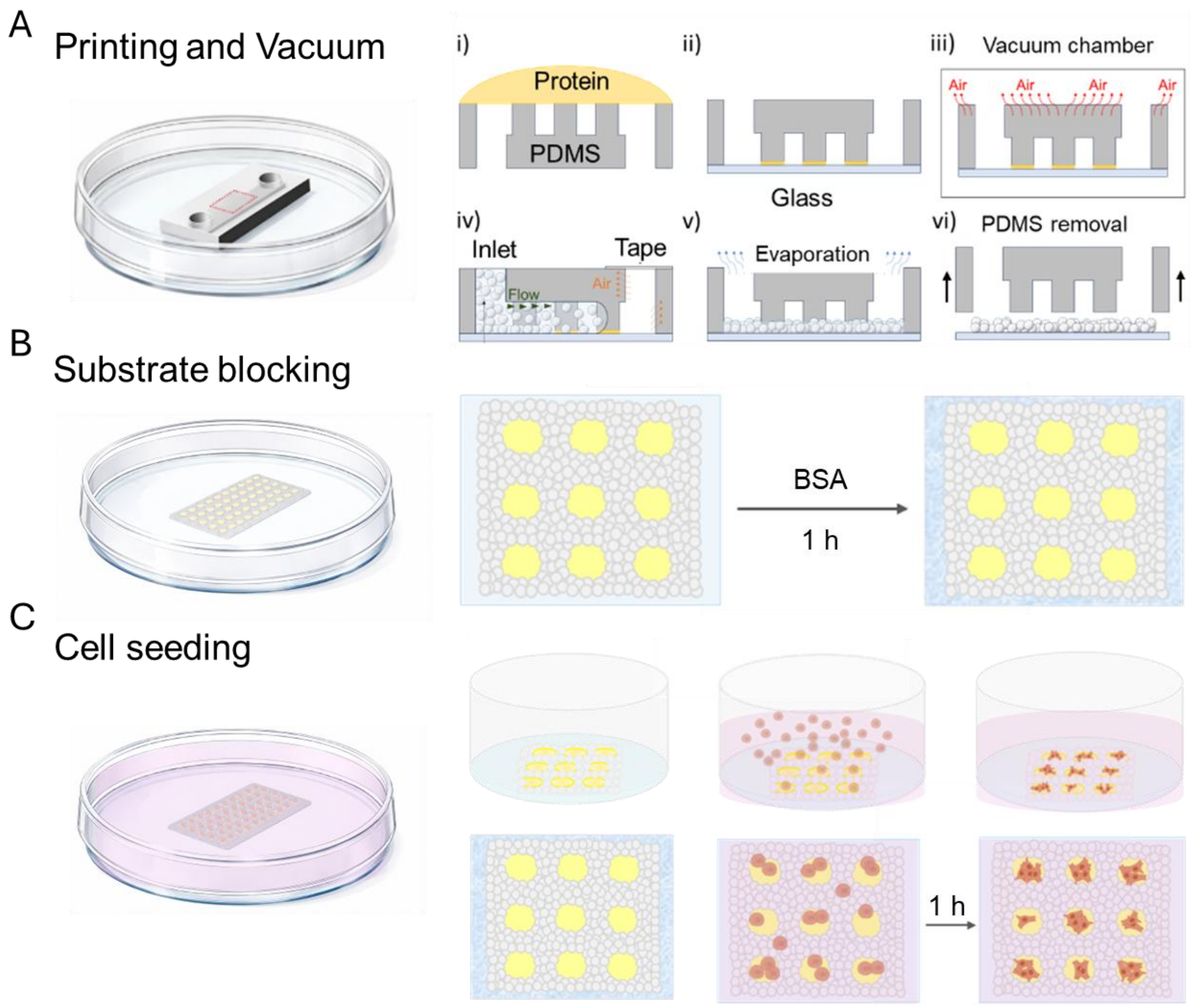


**Figure SI-1. Fabrication of CellStudio substrates trough PnVlitho.** 1) Schematic drawing of microcontact printing: pillars within the PDMS stamp are incubated with a cell adhesion protein solution (i), and after drying, stamp is directly put into contact with the flat surface (ii). PDMS stamps are degassed upon vacuum (iii) that when put into normal atmospheric pressure enables the flow of a microbead suspension between its features (iv). Upon solvent evaporation (v), the microbeads remain bound to the flat surface (vi). B) After PDMS stamp retrieval, CellStudio substrates are incubated with BSA solution acting as blocking agent to avoid unspecific adhesion of cell. C) CellStudio substrates are then incubated with a cell suspension, which results in a multicomponent pattern of small cell clusters surrounded by the microbeads.

**Supporting Information 2: Quantification of IFN-γ secretion**

For quantification and comparison of IFN-γ secretion between activation conditions, fluorescence intensity was converted into the amount of IFN-γ secreted per cell per day. Fluorescence intensity was first measured around each cell cluster using a ring-shaped region of interest (ROI) with an inner diameter of 100 μm and an outer diameter of 110 μm, corresponding to the 10 μm region immediately surrounding each protein dot. The fluorescence intensity measured for each cluster was then converted into a localized IFN-γ concentration (ng $mL^{-1}$) using the calibration curve. To obtain the amount of IFN-γ associated with each cell cluster, a defined local volume was assigned to each patterned spot. Since the CellStudio patterns consisted of 100 μm dots separated by 100 μm, this analytical volume was defined as a hemispherical region with a radius of 100 μm centered on each dot. In this geometry, every point within the outer dome remains within 100 μm of the center of the corresponding cell cluster, while avoiding overlap with the volumes assigned to adjacent patterned spots. The localized concentration was then converted into the total amount of IFN-γ contained within this volume, expressed in fg. Finally, to compare secretion between activation patterns, the amount of IFN-γ associated with each cluster was normalized by the secretion time and by the mean number of cells per dot for each pattern type (P1–P4), yielding IFN-γ secretion in fg $cell^{-1}$ $day^{-1}$ **Figure SI-2**.

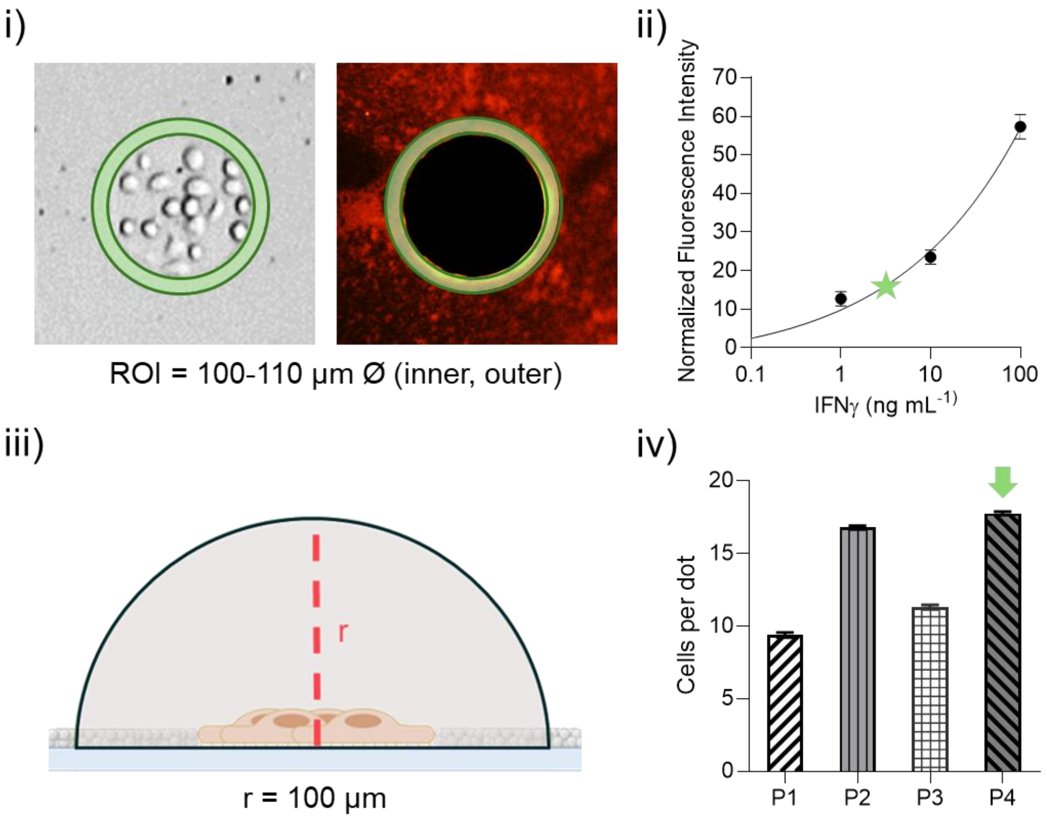


**Figure SI-2. Quantification of IFN-γ secretion.** Fluorescence intensity was measured around each cell cluster using a ring-shaped region of interest (ROI) with an inner diameter of 100 µm and an outer diameter of 110 µm (i). The measured fluorescence intensity was then converted into a localized IFN-γ concentration (ng $mL^{-1}$) using the calibration curve (ii). For each cell cluster, an analytical volume was defined as a hemisphere with a radius of 100 µm centered on the corresponding patterned dot, and this volume was used to convert the concentration value into the amount (fg) of IFN-γ (iii). Finally, to compare secretion between groups, the IFN-γ amount was normalized by the secretion time and by the mean number of cells per dot for each pattern type (P1–P4), yielding IFN-γ secretion per cell per day.
